# NBR1 shuttles between the cytoplasm and nucleus and is essential for nuclear p62 body formation

**DOI:** 10.64898/2026.03.10.710728

**Authors:** Marcus Moe Mauseth, Jana Wurz, Morten Svendsen Næss, Gry Evjen, Hallvard Lauritz Olsvik, Yakubu Princely Abudu, Terje Johansen, Trond Lamark

## Abstract

The selective autophagy receptors SQSTM1/p62 and NBR1 are evolutionary related and involved in autophagy of different substrates including proteins, protein aggregates and organelles. The two proteins interact via their PB1 domains, and essential for their roles in autophagy is the formation of a specific type of condensates named p62 bodies. The scaffold of these structures is formed by the interaction of polymeric p62 with polyubiquitin, but NBR1 is recruited and essential for their formation. Previous studies have shown that p62 contains nuclear export signal (NES) and nuclear localization signal (NLS) motifs and shuttles between the cytoplasm and the nucleus. Its nuclear roles are not fully understood, but there is evidence that p62 is involved in protein quality control in the nucleus. No previous studies have tested if NBR1 is transported into the nucleus. We show here that NBR1 contains two NES motifs and one NLS motif, and like p62, the protein shuttles between the cytoplasm and the nucleus. NBR1 also accumulates in nuclear p62 bodies and the formation of nuclear p62 bodies depends on NBR1.

## INTRODUCTION

SQSTM1/p62 (sequestosome-1) and NBR1 (neighbor of BRCA1 gene 1) are two of the most studied selective autophagy receptors (SARs) in mammals. A characteristic feature of p62 is the formation of so-called p62 bodies (Bjørkøy et al., 2005; Lamark and Johansen, 2021). These are stress-induced cytoplasmic condensates consisting of p62 filaments and polyubiquitinated substrates. Their formation depends on the N-terminal Phox/Bem1p (PB1) and C-terminal Ubiquitin-Associated (UBA) domain in p62 (Lamark et al., 2003; Wilson et al., 2003). PB1 domains are protein-protein interaction domains mediating electrostatic interactions between two PB1-containing proteins (Terasawa et al., 2001). Heteromeric interactions are formed by a specific interaction between the acidic OPCA motif in one PB1 domain and a cluster of basic residues in the other. A few PB1 domain-containing proteins, like p62, are able to self-interact via the PB1 domain. For p62, repeated PB1 mediated “head-to-tail” interactions lead to the formation of long helical filaments (Ciuffa et al., 2015). Polymerization of p62 is essential, and it is the interaction of polymeric p62 with polyubiquitin that induces the formation of p62 bodies (Lamark et al., 2003; Wilson et al., 2003). Their formation is strongly facilitated by an increase in the level of p62, and this at least partially reflects the need for phase separation. The UBA domain of p62 has low basal affinity for ubiquitin (Kirkin et al., 2009), and formation of p62 bodies also strongly depends on post-translational modifications, including phosphorylations and acetylation, that increase the affinity of the UBA domain for ubiquitin (Matsumoto et al., 2011; You et al., 2019).

NBR1 is larger than p62 and contains several additional domains, but the proteins share the same domain architecture including an N-terminal PB1 domain and a C-terminal UBA domain (Lamark et al., 2003). An important difference is that the PB1 domain of NBR1 lacks a basic cluster and therefore does not polymerize like the PB1 domain in p62. Instead, the OPCA motif in the PB1 domain of NBR1 interacts with the basic cluster in the PB1 domain of p62 (Lamark et al., 2003). Microscopy of cytoplasmic p62 bodies indicates that NBR1 is always present in p62 bodies, and the colocalization depends on the PB1 interaction. A single aspartate to glutamate (D50R) mutation in the OPCA motif of NBR1 is sufficient to prevent the localization of NBR1 in p62 bodies (Kirkin et al., 2009). An important role of NBR1 is to facilitate p62 body formation, and siRNA-mediated depletion of NBR1 blocks the formation of endogenous p62 bodies formed in response to the translational inhibitor puromycin (Kirkin et al., 2009). Cytoplasmic condensates of p62 are formed without NBR1 if the level of p62 is very high, but NBR1 facilitates their formation and stress induced formation of endogenous p62 bodies depends on a co-presence of NBR1 (Sanchez-Martin et al., 2020; Zaffagnini et al., 2018). It is not understood why NBR1 is required, and there may be multiple reasons. The number of NBR1 molecules in a cell is normally much lower than the number of p62 molecules, and a sub-stoichiometric concentration of NBR1 relative to p62 is needed for condensate formation (Turco et al., 2021). Binding of NBR1 to a p62 filament caps the basic end of the filament effectively terminating polymerization, and one possible function for NBR1 may be to regulate the length of p62 filaments (Jakobi et al., 2020). Very long filaments are potentially less efficiently packed into dynamic condensates. Other possible roles can be to recruit enzymes or proteins that participate in phase separation, or it may use its UBA domain to increase the binding of p62 filaments to ubiquitin (Turco et al., 2021), or to recruit ubiquitinated substrates.

Autophagy is a cytoplasmic process, but p62 contains nuclear localization signals (NLS) and a nuclear export signal (NES) and the protein shuttles continuously between the nucleus and cytoplasm (Pankiv et al., 2010). The p62 bodies can also be formed in the nucleus, and p62 is efficiently recruited to ubiquitinated aggregates formed in the nucleus. Nuclear p62 bodies are always formed adjacent to PML (promyelocytic leukemia) bodies (Pankiv et al., 2010), and p62 bodies are sometimes engulfed by a PML body (Souquere et al., 2015). A recent study suggests that p62 bodies stabilize PML bodies by sequestering and degrading the E3 ligase RNF4 (Fu et al., 2024). Several studies indicate a role in facilitating proteasomal degradation of nuclear substrates (Fu et al., 2021; Pankiv et al., 2010). Nuclear p62 bodies contain ubiquitinated substrates, the 26S proteasome, ubiquitin conjugating enzymes and deubiquitinating enzymes (DUBs) (Fu et al., 2021; Pankiv et al., 2010). By bringing all these components into a dynamic condensate, p62 bodies are suggested to act as centers for proteasomal degradation of nuclear proteins.

No previous studies have tested if NBR1 can shuttle in and out of the nucleus, or if nuclear p62 bodies contain NBR1. Here we show that NBR1 contains functional NLS and NES motifs and shuttles like p62 between the cytoplasm and the nucleus. NBR1 is shown to accumulate in nuclear p62 bodies, and the formation of nuclear p62 bodies is facilitated strongly by the presence of NBR1.

## RESULTS

### Endogenous NBR1 shuttles between the cytoplasm and the nucleus

NBR1 is a well-established selective autophagy receptor (Rasmussen et al., 2022). However, while NBR1 is predominantly localized to cytoplasm, we observed that a fraction of cells displayed condensed nuclear NBR1 puncta (**Fig. 1A**). This localization pattern is similar to previous observations made for nuclear p62 bodies (Pankiv et al., 2010), and indicates that NBR1 is shuttled between the cytoplasm and the nucleus. To test this, we treated HEK293-, HeLa-, and U2OS cells with leptomycin B (LMB), an inhibitor of exportin-1 (aka CRM1) which mediates the nuclear export of proteins containing a leucine-rich nuclear export signal (NES) (Kudo et al., 1999). Following treatment with LMB, we observed an accumulation of NBR1 in the nucleus with a substantial increase in the number and size of nuclear NBR1 puncta in all three cell lines (**Fig. 1B, -C and -D**).

**Figure 1.**
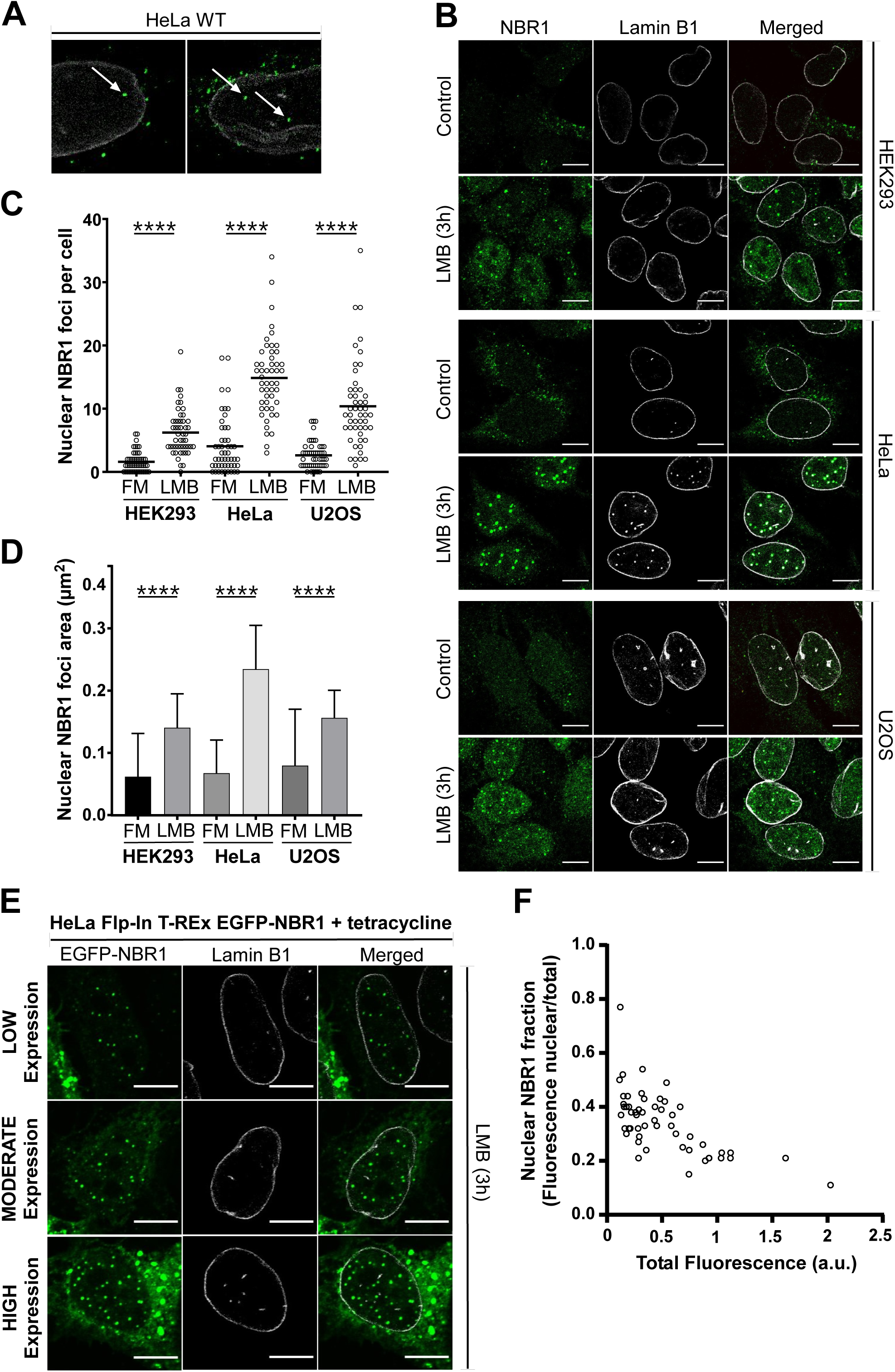
Endogenous NBR1 shuttles between the cytoplasm and the nucleus. (A) HeLa cells stained for endogenous NBR1 (green) and the nuclear envelope marker lamin B1 (white). White arrows indicate nuclear NBR1 puncta. Scale bars equal to 10 µm. (B) HEK293-, HeLa-, and U2OS cells treated with leptomycin B (LMB, 3 h) or not (control). Cells were stained for NBR1 (green) and lamin B1 (white). Scale bars equal to 10 µm. (C) Quantification of number of NBR1 puncta per nucleus in B (n=50 cells). (D) Quantification of mean ± SD area of NBR1 puncta (µm²) in B (n=50 cells). Statistical comparison by One-way ANOVA. ****p < 0.0001. (E) HeLa Flp-In T-REx EGFP-NBR1 cells were treated with tetracycline (24h) to induce expression of EGFP-NBR1 (green) and subsequently subjected to LMB (3h). Cells were stained for lamin B1 (white). (F) Nuclear NBR1 fluorescence in E plotted as a function of total cellular NBR1 fluorescence for individual cells (n=50 cells).

Compared to endogenous NBR1, transiently transfected EGFP-NBR1 has a minor nuclear import after treatment with LMB (**Fig. S1A and -B**). Transiently transfected EGFP-NBR1 is previously shown to accumulate in cytoplasmic aggregates consisting of small vesicles and organelles (Deosaran et al., 2013). To explain the reduced nuclear import observed for transiently transfected NBR1 constructs compared to endogenous NBR1, we hypothesized that overexpression of transfected NBR1 leads to abnormally high protein levels. Since NBR1 self-interacts via its coiled-coil (CC1) domain, excessive expression may promote cytoplasmic aggregation, which in turn sequesters NBR1 and impairs its nuclear import. In comparison, nuclear import of endogenous p62 was seen in cells transfected with EGFP-NBR1 (**Fig. S1A**), although it was noted that some p62 remained in cytoplasmic NBR1 aggregates. In cells transiently transfected with D50R mutated EGFP-NBR1 that does not interact with p62, the import of p62 was more efficient (**Fig. S1A**). This suggests that also the nuclear import of p62 is partially reduced in cells transiently transfected with EGFP-NBR1. To further explore the effect of EGFP-NBR1 overexpression on nuclear import, we made a HeLa Flp-In T-Rex cell line stably expressing EGFP-NBR1 from a tetracycline-induced promoter. By subjecting the cell line to sequential tetracycline- and LMB treatment, we observed a graded EGFP-NBR1 expression level that varied substantially between the cells. Nuclear import upon treatment with LMB also varied between individual cells (**Fig. 1E**). Plotting the fraction of nuclear fluorescence against the total fluorescence revealed that higher expression-levels of NBR1 correlated with a reduced nuclear import upon treatment with LMB (**Fig. 1F**).

### NBR1 accumulates in nuclear p62 bodies

NBR1 and p62 are well-established interaction partners in the cytoplasm and co-localize in cytoplasmic p62 bodies (Bjørkøy et al., 2005; Lamark et al., 2003). To determine if the nuclear NBR1 puncta are p62 bodies, we stained for p62 and proteins known to co-localize with p62 in nuclear bodies (Pankiv et al., 2010). This revealed that the nuclear NBR1 puncta were positive for p62 (>95%), PML (∼70%) and ubiquitinated proteins (FK2) (∼60%) (**Fig. 2A, S1C and - D**). The NBR1 puncta were perfectly co-localized with p62 and ubiquitinated proteins, while PML bodies were adjacent and did not completely overlap with NBR1 puncta (**Fig. S1C**). Previous studies have shown a similar partial co-localization of nuclear p62 with PML (Pankiv et al., 2010). Our conclusion is that nuclear NBR1 accumulates in p62 bodies.

**Figure 2.**
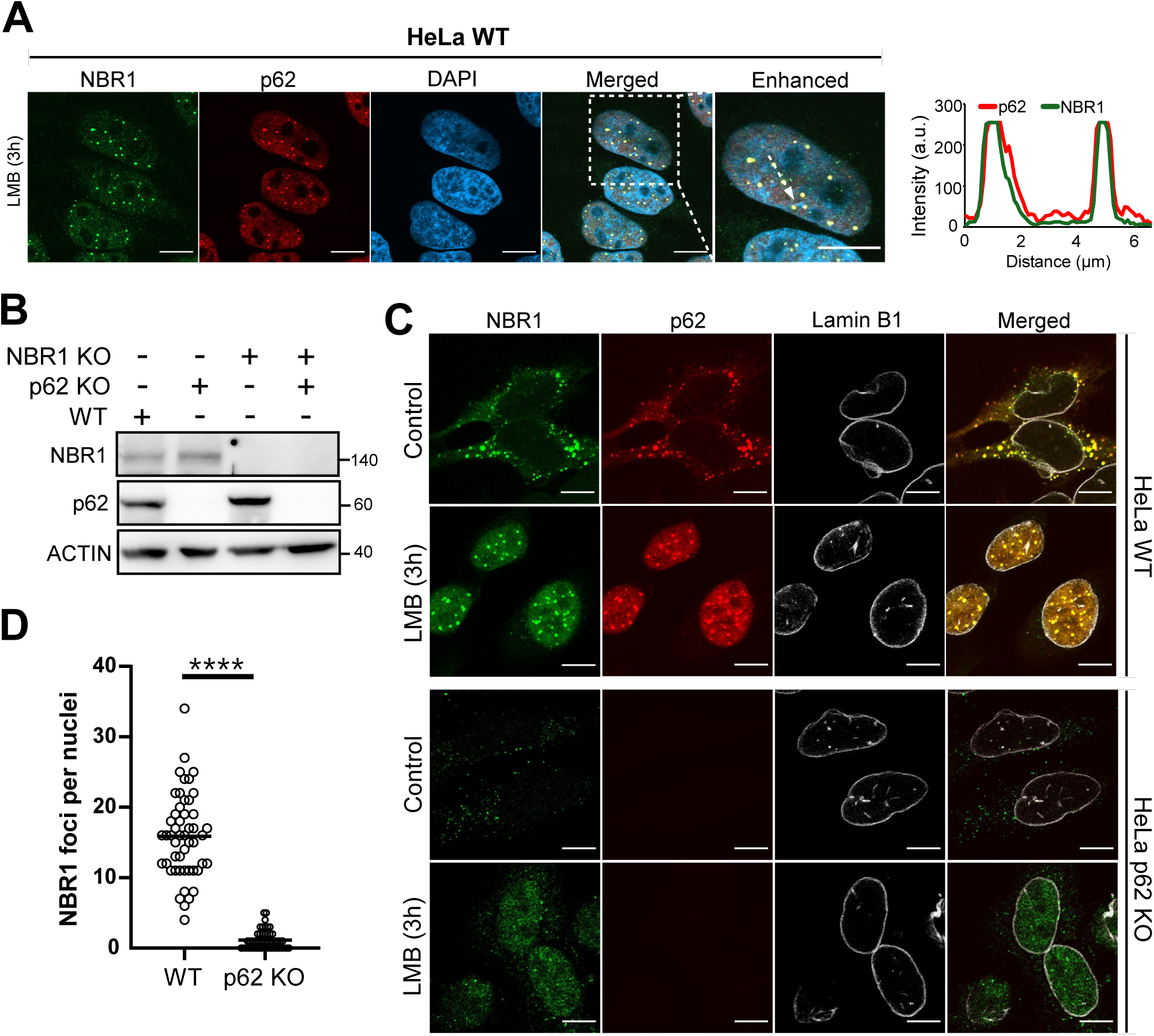
Endogenous NBR1 accumulates in nuclear p62 bodies. (A) HeLa cells were treated with LMB (3 h) and stained for NBR1 (green) and p62 (red). Scale bars equal to 10 µm. Lineplot of co-localization to the right. (B) KO clones were generated from WT HeLa cells. Cell extracts from WT, p62 KO, NBR1 KO, and p62/NBR1 DKO HeLa cells were analyzed by western blotting using the indicated antibodies. (C) WT- and p62 KO HeLa cells were treated or not with LMB (3h). Cells were stained for NBR1 (green), p62 (red) and lamin B1 (white). (D) Quantification of NBR1 foci per nucleus in C (n=50 cells). Statistical comparison by unpaired t-test. ****p < 0.0001.

Having observed the nuclear co-localization of NBR1 with p62, p62 knockout (KO) HeLa cells were made using CRISPR/Cas9 to investigate the effect on nuclear import and nuclear aggregation of NBR1 (**Fig. 2B**). The absence of p62 strongly reduced the number and size of nuclear NBR1 puncta in cells treated with LMB (**Fig. 2C, -D and S1E**). This suggests that the accumulation of endogenous NBR1 in nuclear puncta depends on p62 expression. This is consistent with the conclusion that nuclear NBR1 accumulates in p62 bodies.

### NBR1 contains a nuclear export signal

The accumulation of NBR1 in the nucleus following LMB treatment infers the presence of one or more nuclear export signals (NES). Classical NES motifs follow a distinct pattern, consisting of regularly spaced hydrophobic residues (Φ), interspersed by non-hydrophobic residues. These motifs generally follow the consensus Φ1-(x)2–3-Φ2-(x)2–3-Φ3-x-Φ4, where x denotes any amino acid (Lee et al., 2019). By utilizing the NES-prediction tool LocNES (Xu et al., 2015), we scanned NBR1 for consensus NES motifs (**Fig. 3A**). Two putative NES sequences; NES1 (169-185) and NES2 (557-572) were selected for experimental verification. To assess the functional relevance of these sequences, we generated constructs of NBR1 with NES1 and/or NES2 deleted (**Fig. 3B**). These were transiently transfected into HeLa cells, where we observed nuclear accumulation of EGFP-NBR1 following deletion of NES1 or both NES1 and NES2, but not upon deletion of NES2 alone (**Fig. 3C**). Notably, the combined deletion of NES1 and NES2 resulted in greater nuclear localization than deletion of NES1 alone. These results suggest that NBR1 contains at least two functional NES sequences, with NES1 exhibiting significantly stronger nuclear export activity than NES2 (**Fig. 3D**). To further examine these sequences, we utilized the pRev(1.4)-EGFP reporter system (Henderson and Eleftheriou, 2000). This reporter plasmid encodes a truncated HIV-1 Rev protein fused to EGFP, which localizes exclusively to the nucleus unless complemented by a functional NES. Insertion of candidate sequences into this vector allowed us to monitor NES activity based on EGFP subcellular localization. Transfecting WT HeLa cells with p.Rev(1.4)-GFP, p.Rev(1.4)-NBR1(NES1)-EGFP, p.Rev(1.4)-NBR1(NES2)-EGFP, and p.Rev(1.4)-p62(NES)-EGFP, we observed that only NBR1(NES1) and the positive control, p62(NES) (Pankiv et al., 2010), were able to export the construct from the nucleus (**Fig. 3E and -F**). Taken together, these data strongly indicate that NES1 is the main nuclear export signal in NBR1, although a contributory role for NES2 is also possible.

**Figure 3.**
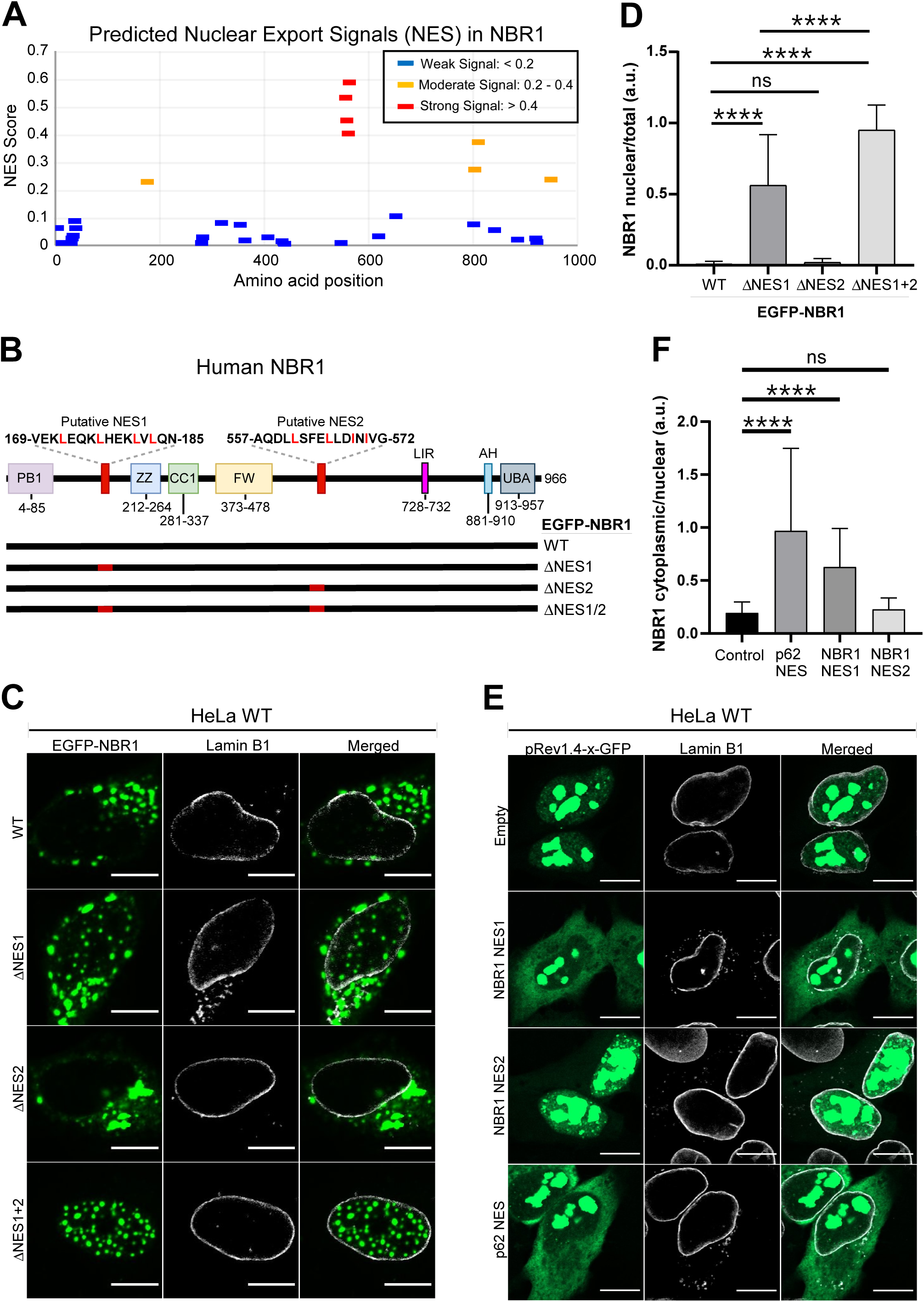
NBR1 contains a putative nuclear export signal. (A) Putative NES signals predicted by LocNES, with the NES score categorized as either strong signal (≥ 0.4), moderate signal (0.2 – 0.4), weak signal (0.1 – 0.2), and lowest signal (< 0.1). (B) Domain architecture of human NBR1, showing localization of two putative NES signals relative to other known domains. At the bottom, NES1 and NES2 deletion constructs are shown with deleted fragments indicated in red. (C) WT HeLa cells were transfected with EGFP-NBR1 (WT), EGFP-NBR1 ΔNES1, EGFP-NBR1 ΔNES2 or EGFP-NBR1 ΔNES1ΔNES2 (ΔNES1/2). Cells were stained for lamin B1 (white). (D) Quantification of nuclear NBR1 in C. Nuclear NBR1 fluorescence of each cell was normalized to the total NBR1 fluorescence in the cell (n=50 cells). Statistical comparison by One-way ANOVA. NS=not significant, ****p < 0.0001. (E) WT HeLa cells transfected with pRev1.4-x-GFP (control without NES) or pRev1.4-x-GFP carrying an insertion of NBR1 NES1 (residues 169-186), NBR1 NES2 (residues 557-574), or p62 NES (residues 303-320). (F) Quantification of cytoplasmic NBR1 fluorescence in E (n=50 cells). The cytoplasmic NBR1 signal of each cell was normalized to the nuclear NBR1 signal in the cell. Statistical comparison by One-way ANOVA. NS=not significant, ****p < 0.0001.

### The coiled-coil 1 (CC1) domain of NBR1 contains a nuclear localization signal

The presence of NES in NBR1 necessitates the existence of nuclear import signal(s) (NLS) or alternative mechanisms capable of facilitating the nuclear localization of NBR1. Classical NLSs can be divided into two categories: monopartite- and bipartite NLSs. Monopartite NLSs typically consist of 4–8 positively charged amino acids, and follows a pattern defined as K (K/R) X (K/R), where X can be any residue. Meanwhile, bipartite NLSs consist of two clusters of 2–3 positively charged amino acids interspersed by a 9–12 amino-acid proline-rich linker region (Lu et al., 2021). However, many NLSs do not follow the amino acid sequence patterns observed in classical NLSs. To address this issue, we created a deletion series of EGFP-NBR1ΔNES1+2 (**Fig. 4A**). Since all constructs were defective in nuclear export, we could then identify the sequence(s) responsible for the nuclear localization of NBR1. A notable feature of transiently transfected EGFP-NBR1ΔNES1/2 constructs is that they accumulate in nuclear puncta, and this was also seen for EGFP-NBR1Δ1-185 that cannot interact with p62 (**Fig. 4B**). We showed above that endogenous NBR1 does not readily form nuclear puncta in cells lacking p62 (**Fig. 2C**), but a few small nuclear puncta could be seen in these cells. Transient transfection results in much higher levels of nuclear NBR1, and this presumably explains the observed p62 independent nuclear aggregation seen for EGFP-NBR1 Δ1-185. The endogenous level of NBR1 is presumably too low for this to occur and NBR1 is probably rapidly exported out from the nucleus unless it accumulates with p62 in nuclear p62 bodies.

**Figure 4.**
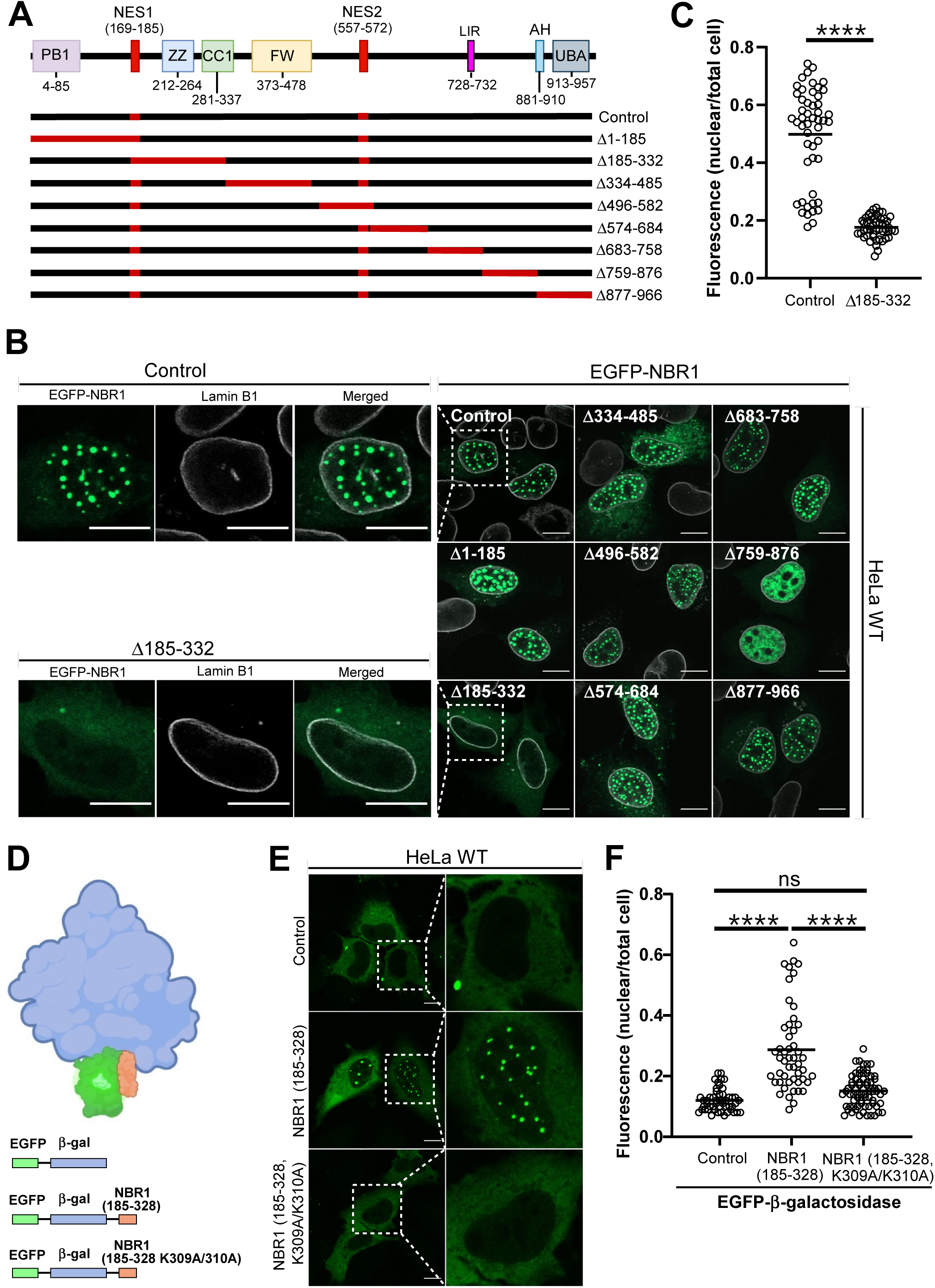
NBR1 contains a putative nuclear import signal. (A) Serial deletions of EGFP-NBR1, with deleted fragments indicated in red. All NBR1 constructs carry a deletion of NES1 and NES2. (B) EGFP-NBR1 deletion constructs were transfected in WT HeLa cells. Cells were stained for lamin B1 (white) and show NBR1 in green. At left, enlarged view of control and Δ185-332. Scale bars equal to 10 µm. (C) Quantification of data in B for control and Δ185-332. Nuclear NBR1 fluorescence normalized to whole-cell fluorescence (n = 50 cells). Unpaired t-test used for statistical comparison; ****p < 0.0001. (D) At top, schematic illustration of EGFP-β-galactosidase-NBR1 (185-328), indicating relative sizes of EGFP, β-galactosidase and NBR1 (185-328). At bottom, domain architecture of fusion proteins expressed in E. (E) WT HeLa cells transfected with the indicated EGFP-β-galactosidase constructs. (F) Quantification of nuclear NBR1 fluorescence of E (n = 50 cells). Nuclear fluorescence was normalized to total cell fluorescence. One-way ANOVA used for statistical comparison; NS=not significant, ****p < 0.0001.

Transient transfection revealed that a deletion of amino acid residues 185-332 including the ZZ and CC1 domains (construct 3), resulted in cytoplasmic localization (**Fig. 4B and -C**). All other deletion constructs accumulated in the nucleus (**Fig. 4B**). Using the prediction tool NLStradamus, which employs hidden Markov models (HMMs) to identify novel NLSs (Nguyen Ba et al., 2009), the highest probability for an NLS in NBR1 was predicted for residues 300 to 328 in the CC1 domain of NBR1. This fits well with our observations and indicates the presence of at least one NLS in the CC1 domain of NBR1 (residues 281-337). We verified this by fusing NBR1(185-328) to EGFP-β-galactosidase (**Fig. 4D**). The large size of β-galactosidase prevents nuclear import of EGFP-β-galactosidase by diffusion, and we previously used the same construct for the identification of NLS sequences in p62 (Pankiv et al., 2010). Transfection of EGFP-β-galactosidase-NBR1(185-328) resulted in nuclear localization (**Fig. 4E**), and this verified the existence of at least one functional NLS in this NBR1 fragment. The mutation of two specific lysine residues in the CC1 domain (K309A and K310A), ablated the nuclear localization of EGFP-β-galactosidase-NBR1(185-328) (**Fig. 4E and -F**). The effect of this mutation strongly indicates that an NLS is located in the CC1 domain, and most likely includes the two mutated residues. Our data does not exclude the existence of other weaker NLS sequences in NBR1, but no import was seen when testing other predicted candidates in the β-galactosidase vector.

### NBR1 is essential for formation of nuclear p62 bodies

One possible nuclear role for NBR1 could be to facilitate nuclear p62 body formation. To assess this, we knocked out NBR1 in WT HeLa cells using CRISPR/Cas9 (**Fig. 2B**) and subsequently treated WT- and NBR1 KO HeLa cells with LMB for 3h. Following LMB treatment, we observed a significant reduction in nuclear p62 bodies in the NBR1 KO cells compared to the WT cells (**Fig. 5A, -B and -C**). The formation of nuclear p62 bodies is under normal conditions strongly limited by the lack of nuclear accumulation of these two proteins. In contrast to this, we showed above that NES deleted EGFP-NBR1 is not exported and accumulates in nuclear puncta (**Fig. 3C**). To test if a nuclear accumulation of NBR1 alone is sufficient to affect the localization pattern of p62, HeLa cells were transiently transfected with EGFP-NBR1ΔNES1/2 and endogenous p62 detected by immunostaining. It appeared that endogenous p62 accumulated in the nucleus of cells expressing NES-deleted NBR1, even in the absence of LMB-mediated inhibition of nuclear export (**Fig. 5D**). This indicates that shuttling p62 is entrapped by nuclear NBR1, preventing its return to the cytoplasm. Next, we tested if the expression of a D50R mutated NBR1 construct has the same effect. The direct interaction between NBR1 and p62 depends on their PB1 domains and is abolished by a D50R mutation in the PB1 domain in NBR1 (Lamark et al., 2003). EGFP-NBR1 D50R ΔNES1ΔNES2 formed nuclear puncta despite of its inability to interact with p62, but there was no visible nuclear accumulation of endogenous p62 and no p62 was seen in nuclear NBR1 puncta (**Fig. 5D**). Our conclusion is therefore that a direct interaction of NBR1 with p62 facilitates the formation of nuclear p62 bodies.

**Figure 5.**
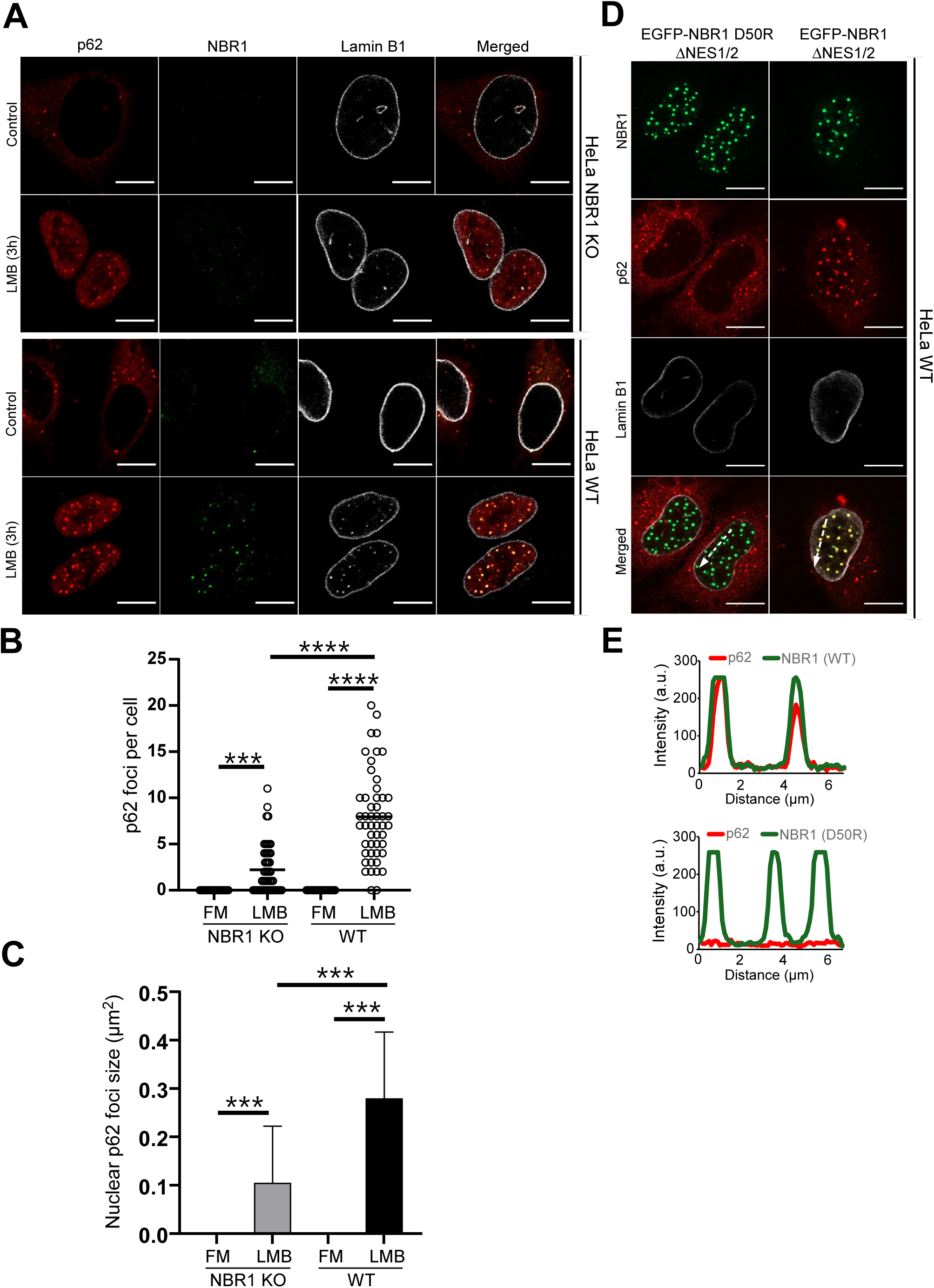
NBR1 is required for efficient formation of nuclear p62 bodies. (A) NBR1 KO-or WT HeLa cells were treated or not with LMB (3h). Cells were stained for p62 (red), NBR1 (green) and lamin B1 (white). (B) Quantification of nuclear p62 puncta in A (n=50 cells). Statistical comparison by One-way ANOVA. ****p < 0.0001, ***p < 0.001. (C) Quantification of mean ± SD area of p62 puncta (µm²) in A (n=50 cells). Statistical comparison by One-way ANOVA. ****p < 0.0001. (D) WT HeLa cells transfected with NES deleted EGFP-NBR1 D50R or NES deleted EGFP-NBR1. Cells were stained for p62 (red) and lamin B1 (white). (E) Lineplots of co-localization of indicated puncta from the merged images in (D).

### NBR1 and p62 are mutually dependent on each other for effective recruitment to nuclear Ataxin-1 inclusions and promote proteasome recruitment to the inclusion surface

An important question is whether NBR1 is also essential for the effective recruitment of p62 to nuclear protein inclusions. Using Hela or HEK293 cells, we previously reported that p62 is recruited to nuclear inclusions generated by mutant Ataxin1 (Atx84Q) and facilitates the recruitment of proteasomes to these structures (Pankiv et al., 2010). To test whether this process is contingent on NBR1, we co-transfected WT-, NBR1 KO-, and p62 KO cells with EGFP-Atx84Q. This pathogenic Ataxin-1 variant contains a polyglutamine expansion of 84 residues, which is the main etiological factor of spinocerebellar ataxia type 1 (Cummings et al., 1998). By transfecting WT HeLa cells with EGFP-Atx84Q, we observed the formation of spherical nuclear protein inclusions in all three cell lines (**Fig. 6A**). In WT cells treated with LMB to increase the nuclear level of endogenous p62 and NBR1, immunostaining revealed a strong recruitment of NBR1 and p62 positive puncta at the surface of larger Atx84Q inclusions (**Fig. 6A and -B**). In contrast, p62 recruitment to Ataxin-1 inclusions was significantly reduced in NBR1 KO cells, and NBR1 recruitment was similarly diminished in p62 KO cells (**Fig. 6A and -B**). These findings indicate a mutual dependency of NBR1 and p62 for efficient recruitment to Atx84Q inclusions. WT cells transfected with EGFP-Atx84Q were also stained for the proteasomal subunit β2. A strong co-localization of this proteasomal subunit in puncta containing endogenous NBR1 indicated a co-recruitment of proteasomes with p62 and NBR1 into Atx84Q inclusions (**Fig. 6C**). No specific co-localization of the proteasomal subunit β2 with Atx84Q was seen in p62/NBR1 double KO (DKO) cells (**Fig. 6C**). Our data therefore supports a role for p62 and NBR1 in bringing proteasomes to nuclear inclusions. Co-transfection of EGFP-Atx84Q with NES-deleted p62 and NBR1 constructs confirmed that nuclear puncta of p62 and NBR1 are formed adjacent to Ataxin1 inclusions (**Fig. 6D**).

**Figure 6.**
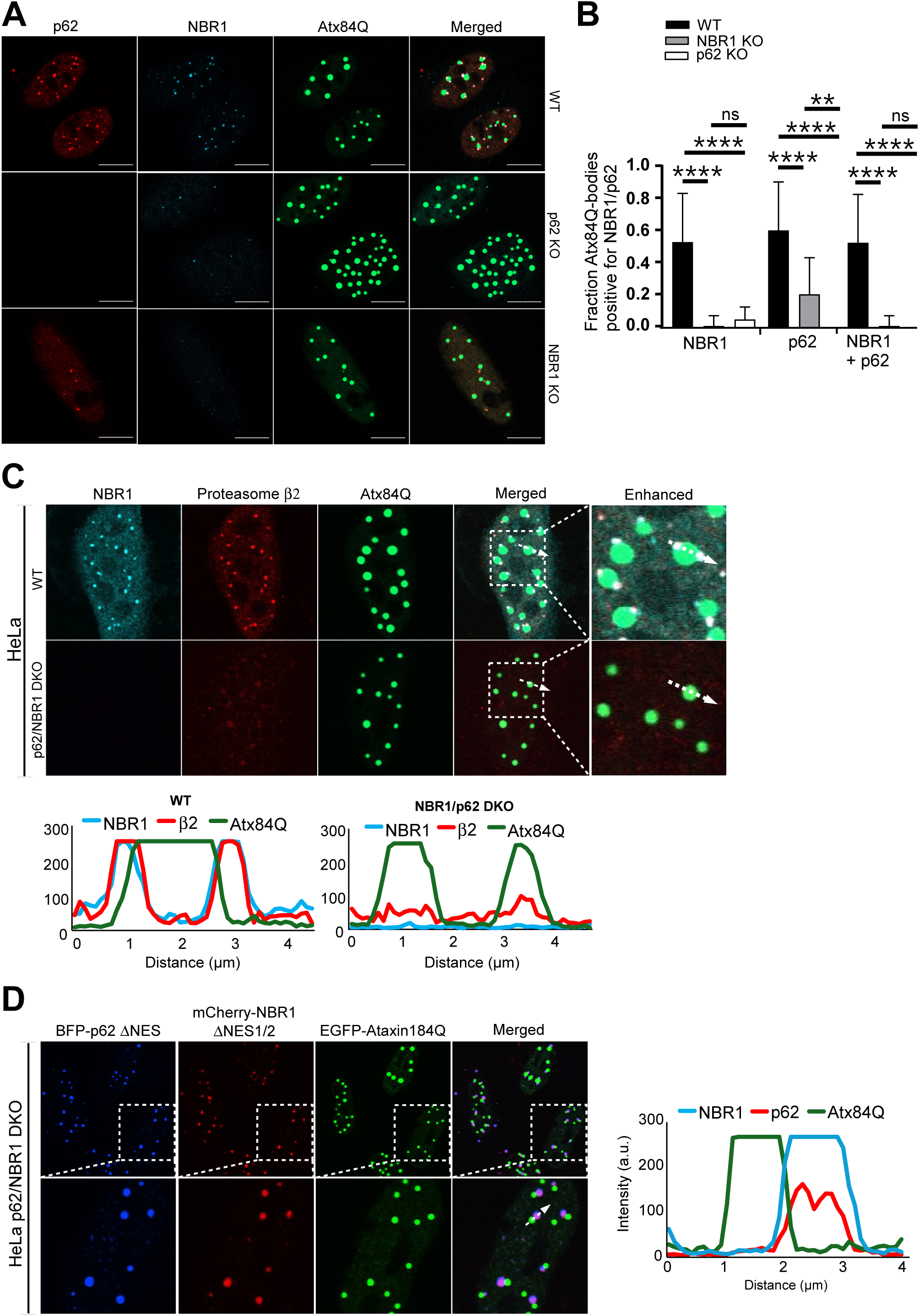
NBR1 and p62 are mutually dependent on each other for effective recruitment to nuclear Ataxin-1 inclusions and promote proteasome recruitment to the inclusion surface. (A) WT-, p62 KO-, and NBR1 KO HeLa cells transfected with EGFP-Atx84Q (green). 20 hours after transfection, cells were treated with LMB (3 hours) and stained for endogenous p62 (red) and endogenous NBR1 (blue). (B) Quantification of Atx84Q inclusions positive for p62 and/or NBR1 in E (n = 50 cells). Two-way ANOVA was used for statistical comparison. NS = not significant, ****p < 0.0001, **p < 0.01. (C) WT- and p62/NBR1 DKO HeLa cells were transfected with EGFP-Atx84Q (green). Twenty hours after transfection, cells were treated with LMB (3 hours) and stained for endogenous NBR1 (blue) and endogenous proteasomal β2 subunit (red). Lineplots of co-localization of indicated puncta are shown below the images. (D) p62/NBR1 DKO HeLa cells were transfected with BFP-p62ΔNES (blue), mCherry-NBR1ΔNES1/2 (red) and EGFP-Atx84Q (green). To the left, lineplot of co-localization of indicated puncta.

### NBR1 changes the dynamics of nuclear condensation of p62

Next, we examined the dynamics of nuclear p62 body formation. FRAP analyses of EGFP positive puncta were performed to test if nuclear p62 bodies have liquid phase properties. To secure an efficient nuclear accumulation, we transiently transfected NES deleted constructs of p62 and NBR1 into HeLa p62/NBR1 DKO cells. When transiently expressing NES deleted constructs, the nuclear expression is high, and all combinations resulted in the formation of nuclear aggregates. Since the recovery rate may depend on the size of nuclear puncta (**Fig. S2**), we selected EGFP-p62ΔNES puncta of similar size. After bleaching, an efficient recovery of EGFP-p62ΔNES puncta was seen both if expressed alone or together with mCherry-NBR1ΔNES1/2 (**Fig. 7A-C**). This indicated that nuclear p62 bodies have liquid phase properties, both in the presence or absence of co-expressed NBR1. To further investigate the dynamics of nuclear p62 body formation, p62/NBR1 DKO HeLa cells transfected with EGFP-p62ΔNES alone or together with mCherry-NBR1ΔNES1/2, were subjected to treatment with 1,6-hexanediol. 1,6-hexanediol disrupts weak hydrophobic protein-protein or protein-RNA interactions of liquid condensates but does not interfere with solid protein aggregates (Kroschwald et al., 2017). Treatment with 1,6-hexanediol resulted in a rapid loss of p62 bodies, and this supported the conclusion that they are liquid condensates (**Fig. 7D**). Using 1,6-hexanediol, we observed that the recovery of EGFP-p62ΔNES puncta was faster if mCherry-NBR1ΔNES1/2 was co-expressed (**Fig 7D**). Nuclear puncta formed by EGFP-NBR1ΔNES1/2 alone were also disrupted by the treatment with 1,6-hexanediol, indicating that their formation at least partially depends on liquid phase condensation. However, nuclear puncta formed by EGFP-NBR1 alone did not recover noticeably in the measured timespan (**Fig 6D**). Taking all our data together, we conclude that NBR1 facilitates efficient formation of nuclear p62 bodies.

**Figure 7.**
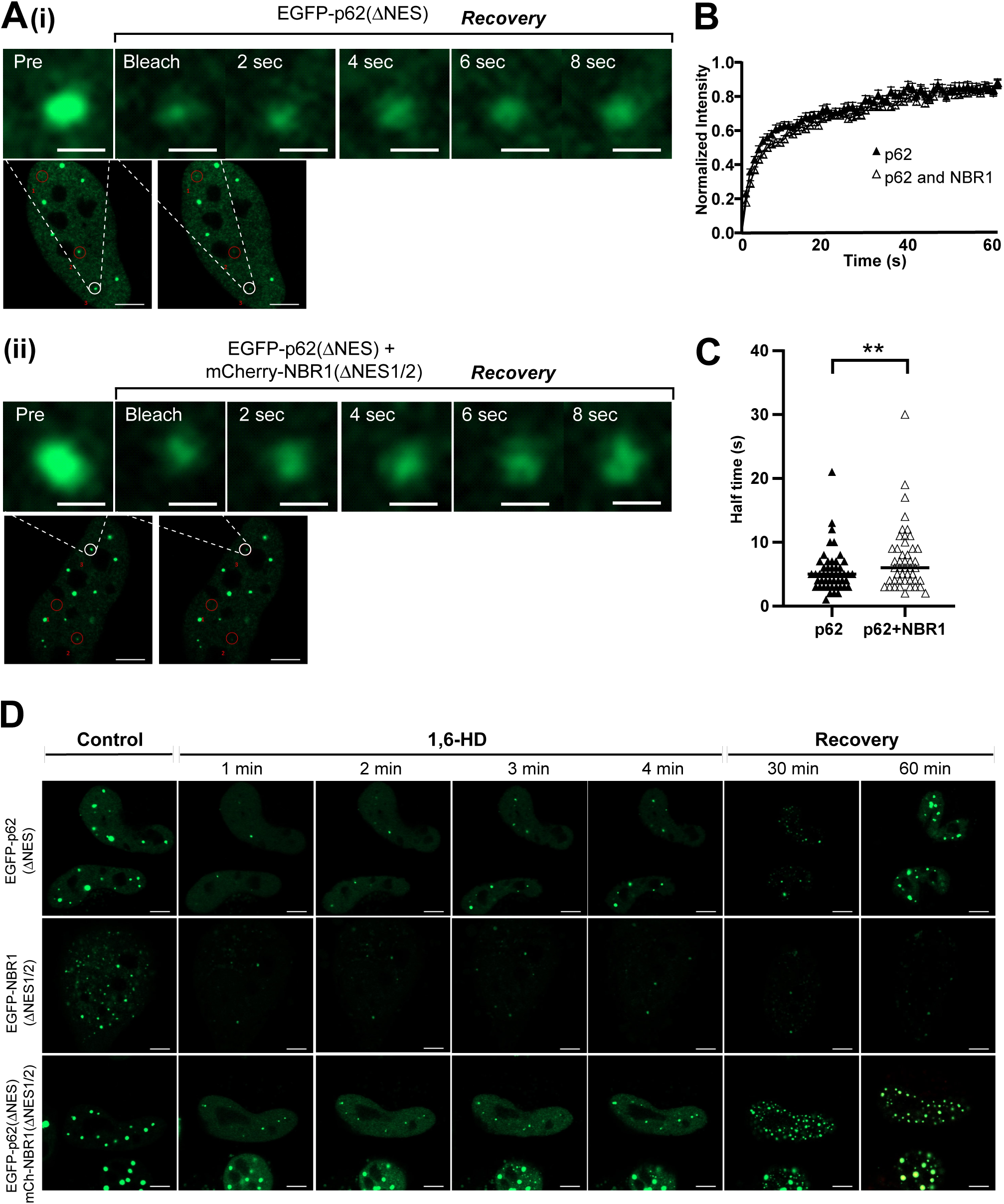
The dynamics of p62 nuclear bodies are influenced by NBR1. (A) FRAP analysis of EGFP positive nuclear puncta in WT HeLa cells transfected with EGFP-p62ΔNES alone (i) or co-transfected with EGFP-p62ΔNES and mCherry-NBR1ΔNES1/2 (ii). (B) FRAP analyses of EGFP positive nuclear puncta in HeLa p62/NBR1 DKO cells transfected with EGFP-p62ΔNES, either alone or together with Cherry-NBR1ΔNES1/2. Intensity values of EGFP-p62 dots with NBR1 co-transfection (n=46) or without (n=49) were measured over time in three separate experiments (n=49 for p62 alone, and 46 for p62 with NBR1 co-expression). Values were normalized to pre-bleach intensity and corrected for photobleaching. Mean ± SEM values are plotted. (C) Recovery half-time (t_1/2_) values for EGFP-p62 puncta analyzed with FRAP. HeLa p62/NBR1 DKO cells were transfected with EGFP-p62 alone or together with mCherry-NBR1ΔNES1/2. Statistical comparison by t-test with Welch correction; p<0.05. (D) Time-lapse of condensate disruption and recovery following 1,6-HD treatment. HeLa p62/NBR1 DKO cells were transfected with EGFP-p62ΔNES alone, EGFP-NBR1ΔNES1/2 alone, or EGFP-p62ΔNES and mCherry-NBR1ΔNES1/2 together. The depicted experiment represents one out of three independent experiments. Scale bars are equal to 10 µm.

## DISCUSSION

We reported previously that p62/SQSTM1 is recruited to nuclear inclusions of polyglutamine-expanded Ataxin1Q84 and facilitates the recruitment of proteasomes to these structures (Pankiv et al., 2010). Furthermore, the proteolytic role of p62 in the nucleus has been further established by confirming the degradation of c-myc in p62 condensates (Fu et al., 2021). Nuclear p62 inclusions may have additional pathological consequences, since it has been found in nuclear inclusions of a murine Huntington disease model (Nagaoka et al., 2004), and in patients with frontotemporal lobar degeneration with ubiquitin-positive inclusions (Pikkarainen et al., 2008). However, while these findings infer a potentially critical role of p62 in the nucleus, little is known regarding the nuclear function of NBR1. We show here that NBR1 is essential for the formation of p62 bodies in the nucleus. Based on microscopy of endogenous and ectopically expressed nuclear p62 bodies, our conclusion is that all nuclear p62 bodies contain NBR1. Our data also indicate that the formation of nuclear p62 bodies is very inefficient in cells depleted of NBR1.

As previously seen for p62 (Pankiv et al., 2010), our data shows that nuclear shuttling of NBR1 is driven by NLS and NES signals. One functional NLS signal was identified in the CC1 domain of NBR1. The localization of an NLS in a coiled-coil domain was unexpected, but previous studies have shown that an NLS motif may function as part of a coiled-coil domain i.e. in the human transcription factors STAT3, STAT5 and HSF2 (Ma et al., 2003; Sheldon and Kingston, 1993; Shin and Reich, 2013). What should be noted is that the CC1 domain of NBR1 is responsible for self-interaction of NBR1. The PB1 domain of NBR1 is monomeric, but NBR1 can use its CC1 domain to self-interact (Kirkin et al., 2009). An important question is therefore whether the NLS is exposed on the surface of the helix in monomeric NBR1 and likewise if it is exposed on the surface of dimerized/oligomerized NBR1. It also remains to be tested if nuclear import of NBR1 is regulated by CC1-mediated self-interaction.

We also identified two functional NES motifs in NBR1. NES1 (residues 168-185) is the main NES in NBR1 and is located N-terminal to the ZZ domain. NES2 (residues 561-573) partially overlaps with the LC3/GABARAP interacting LIR2 motif of NBR1. Although NES1 was found to be the strongest NES in NBR1, our data strongly suggest that NES2 participates in nuclear export of NBR1. Deletion of both NES motifs gave a complete nuclear localization of NBR1. This shows that NES signals are needed to maintain a cytoplasmic pool of NBR1. Presence of two NES motifs and that mutation of both are needed to give complete nuclear localization of the protein in question has been reported for CPEB1 (Cytoplasmic polyadenylation element-binding protein 1) (Ernoult-Lange et al., 2009), the protein kinase MST1/STK4 (Ura et al., 2001), and the tumor suppressor protein APC (Neufeld et al., 2000).

The conclusion that NBR1 is essential for the formation of nuclear p62 bodies infers that further studies of functions of nuclear p62 bodies should include NBR1. In the cytoplasm, NBR1 is known to bind to membranes and overexpressed NBR1 accumulates in vesicle clusters (Deosaran et al., 2013; Mardakheh et al., 2010). In the nucleus, there are no vesicles. We therefore expected a diffuse nuclear localization pattern of nuclear NBR1 in cells lacking p62, but the protein partially accumulated in protein aggregates. We consider these NBR1 aggregates formed by NBR1 without p62 to be a result of unspecific aggregation caused by a high protein concentration. Under normal conditions, NBR1 will more easily form condensates with p62 and the formation of NBR1 specific protein aggregates is unlikely. It is more interesting to consider possible roles displayed by diffuse fractions of p62 or NBR1, and these proteins may have independent roles that may or may not be related to the formation of p62 bodies. Our data show that nuclear p62 and NBR1 often have a diffuse localization pattern. It is possible that the diffuse fraction contains small condensates that are not detected as visible puncta in the light microscope. Individual roles of diffuse fractions of p62 and NBR1 may be to identify substrates that are eventually recruited into nuclear p62 bodies. Hence, an important nuclear role for NBR1 is presumably to identify nuclear substrates for degradation by the proteasome, and this may include substrates that are not efficiently recruited by p62.

We have shown previously that NBR1 must bind directly to p62 via its PB1 domain to be in p62 bodies (Kirkin et al., 2009). By expressing D50R mutated NBR1, we show here that this is true also for nuclear p62 bodies. It is likely that an important role for NBR1 is to recruit ubiquitinated substrates to the p62 body.

## MATERIALS AND METHODS

### Plasmids

Conventional cloning was done using restriction enzymes and DNA ligase from New England Biolabs. Gateway LR recombination reactions were done using the Gateway cloning system (Invitrogen) and as described in the Gateway cloning technology instruction manual. In-Fusion cloning was done using the In-Fusion cloning kit by Takara and oligonucleotides suggested by Takara or SnapGene software (www.snapgene.com). Larger deletions were introduced with In-Fusion cloning in accordance with tabulated guidelines (Takara). Point mutants were made using the QuikChange site-directed mutagenesis kit (Stratagene). Verification of generated plasmid constructs was done by restriction digestion and/or DNA sequencing (BigDye; Applied Biosystems). Oligonucleotides for PCR, site-directed mutagenesis, or DNA sequencing were ordered from ThermoFisher. Further details pertaining to the construction of plasmids are available upon request.

Gateway destination vectors pDest-EGFP-C1 (Lamark et al., 2003), and pDest-mCherry-C1 (Kirkin et al., 2009), and Gateway entry clones pENTR-NBR1 and pENTR-NBR1 D5OR are previously described (Lamark et al., 2003). pEGFP-C1-Ataxin1Q84 (conventional plasmid for expressing AtaxinQ84 fused to EGFP) (Pankiv et al., 2010), pDest-EGFP-p62Δ321-349 (Gateway expression clone for expression of NES deleted p62 fused to EGFP) (Pankiv et al., 2010), pcDNA5/FRT/TO-EGFP-NBR1 (plasmid for stable expression of EGFP-tagged NBR1 in mammalian cells containing a Flp-In cassette) (Larsen et al., 2010), are also previously described.

β-galactosidase fusions: pEGFP-βGal-N1 was previously made (Pankiv et al., 2010), and carries β-galactosidase from *E-coli* cloned into the pEGFP-N1 vector (Invitrogen). The resulting β-Gal-EGFP fusion construct is not imported into the nucleus of mammalian cells, and fragments of NBR1 were subcloned into this vector to test if they have nuclear import signals. pEGFP-βGal-N1-NBR1(185-328) was made by inserting NBR1(185-328) into pEGFP-βGal-N1. The vector was first linearized with *Hind*III and *Xma*I, and NBR1(185-328) produced as a PCR product from pENTR-NBR1, using primers 5-GATCTCGAGCAAGCTAACCCATCCTTGGGTTCTTGT-3 (forward) and 5-CATTGATCCACCCGGCCACAGGTGAATTTTCCTATGGAGT-3 (reverse). The PCR product was then mixed with linearized pEGFP-βGal-N1 and inserted by In-Fusion cloning. pEGFP-βGal-N1-NBR1(185-328) KK309/310AA was made by site-directed mutagenesis of pEGFP-βGal-N1-NBR1(185-328).

Gateway entry clones pENTR-NBR1 Δ1-185, pENTR-NBR1 D50R Δ185-332, pENTR-NBR1 D50R Δ334-485, pENTR-NBR1 D50R Δ496-582, pENTR-NBR1 D50R Δ574-684, pENTR-NBR1 D50R Δ683-758, pENTR-NBR1 D50R Δ759-876, and pENTR-NBR1 D50R Δ877-966 were made by conventional cloning by deleting the indicated residues from pENTR-NBR1 D50R. Gateway destination clones pDest-EGFP-NBR1 Δ1-185, pDest-EGFP-NBR1 D50R Δ185-332, pDest-EGFP-NBR1 D50R Δ334-485, pDest-EGFP-NBR1 D50R Δ496-582, pDest-EGFP-NBR1 D50R Δ574-684, pDest-EGFP-NBR1 D50R Δ683-758, pDest-EGFP-NBR1 D50R Δ759-876, and pDest-EGFP-NBR1 D50R Δ877-966 were made by gateway LR reactions where the above entry clones were mixed with pDest-EGFP-C1.

Gateway destination clones carrying deletions of NES1 and/or NES2, i.e. pDest-EGFP-NBR1ΔNES1, pDest-EGFP-NBR1ΔNES2, pDest-EGFP-NBR1ΔNES1ΔNES2, pDest-EGFP-NBR1 D50R ΔNES1ΔNES2, pDest-EGFP-NBR1 Δ1-185 ΔNES2, pDest-EGFP-NBR1 D50R Δ185-332 ΔNES1ΔNES2, pDest-EGFP-NBR1 D50R Δ334-485 ΔNES1ΔNES2, pDest-EGFP-NBR1 D50R Δ496-582 ΔNES1, pDest-EGFP-NBR1 D50R Δ574-684 ΔNES1ΔNES2, pDest-EGFP-NBR1 D50R Δ683-758 ΔNES1ΔNES2, pDest-EGFP-NBR1 D50R Δ759-876 ΔNES1ΔNES2, pDest-EGFP-NBR1 D50R Δ877-966 ΔNES1ΔNES2, and pDest-mCherry-NBR1 ΔNES1ΔNES2, were all made from corresponding plasmids lacking NES deletions. The NES deletions were made by In-Fusion cloning, using primers 5-TTAACGAACCATCCTTGGGTTCTTGTCCC-3 (forward) and 5-AGGATGGTTCGTTAACCACTTGTTCTCTGAAC-3 (reverse) for deletion of NES1 (Δ169-185; ΔTVEKLEQKLHEKLVLQN), and primers 5-TTCTGACTTTGGAGAGAGTGCCCCACAAC-3 (forward) and 5-TCTCCAAAGTCAGAAGATCCACAGATGGGATG-3 (reverse) for deletion of NES2 (Δ557-572; ΔAQDLLSFELLDINIVQE). Exceptions were that primers 5-TTAACGAAAACAATTCAATCCATGGACTCCAG-3 (forward) and 5-AATTGTTTTCGTTAACCACTTGTTCTCTG-3 (reverse) were used for deleting NES1 (Δ169-185) from pENTR NBR1 D50R Δ185-332, and primers 5-CCCAGGACGAGCTTGGAAATGAGAAGGAGG-3 (forward) and 5-CAAGCTCGTCCTGGGCAGTCAGAAGATCC-3 (reverse) for deleting NES2 (Δ560-571; ΔLLSFELLDINIVQ) from pENTR NBR1_SR2_ D50R Δ574-684.

pRev1.4-GFP (Henderson and Eleftheriou, 2000), and pRev1.4-p62(303-320)-EGFP (Pankiv et al., 2010), were previously made. pRev1.4 is a NES deficient mutant of HIV-1 Rev1.4 protein, and a nuclear localization signal in Rev1.4 is responsible for nuclear import of pRev1.4-EGFP. pRev1.4-NBR1(169-186)-EGFP and pRev1.4-NBR1(557-574)-EGFP were made by inserting NBR1-NES1 (residues 169-186) or NBR1-NES2 (residues 557-574) between the nuclear Rev1.4 protein and EGFP. NES1 was inserted by hybridization of oligonucleotides 5-GATCCAACGGTTGAGAAGCTTGAACAGAAATTACATGAAAAGCTTGTCCTCCAGA ACCCACCA-3 and 5-CCGGTGGTGGGTTCTGGAGGACAAGCTTTTCATGTAATTTCTGTTCAAGCTTCTCA ACCGTTG-3, followed by cloning into pRev1.4-GFP linearized with *Bam*HI and *Age*I. NES2 was similarly inserted into *Bam*HI and *Age*I cut vector, but the hybridized oligonucleotides were GATCCAGCCCAGGACCTGCTGTCCTTTGAGCTGTTGGATATAAACATTGTTCAAGAGTTGCCA-3 and 5-CCGGTGGCAACTCTTGAACAATGTTTATATCCAACAGCTCAAAGGACAGCAGGTCCTGGGCTG-3.

### Antibodies

The following primary antibodies were used for immunostaining of cells: anti-p62/SQSTM1 (C-terminus) guinea pig polyclonal (Progen #GP62-C), anti-NBR1 mouse monoclonal (Santa Cruz Biotechnology #sc-130380), anti-PML rabbit monoclonal (Abcam #ab179466), anti-FK2 mouse monoclonal (Enzo Life Sciences #BML-PW8810), anti-lamin B1 rabbit polyclonal (Abcam #ab16048), anti-20S proteasome β2 mouse monoclonal (Santa Cruz Biotechnology #sc-515066). The following primary antibodies were used for western blots: anti-p62-Lck ligand mouse monoclonal (BD Biosciences #610833), anti-NBR1 mouse monoclonal (Santa Cruz Biotechnology #sc-130380), anti-Actin rabbit polyclonal (Sigma-Aldrich #A2066). The following secondary antibodies were used: goat anti-mouse IgG, Alexa Fluor™ 488 (Invitrogen #A-11001); goat anti-guinea pig IgG, Alexa Fluor™ 555 (Invitrogen #A-21435), goat anti-rabbit IgG, Alexa Fluor™ 647 (Invitrogen #A-21245), goat anti-mouse IgG, HRP-conjugated (BD Biosciences #554002), goat anti-rabbit IgG, HRP-conjugated (BD Biosciences #554021).

### Reagents

The following reagents were used: 1,6-hexanediol (Sigma-Aldrich #240117), puromycin (Sigma-Aldrich, P8833), DMEM (Sigma-Aldrich #D6429), Penicillin-Streptomycin (Sigma-Aldrich #P4333), FBS (Sigma-Aldrich, #F7524), hygromycin (Thermo Fisher Scientific #10687010), blasticidin (Thermo Fisher Scientific #A1113903), TransIT®-LT1 Transfection Reagent (Mirus #MIR 2304), Leptomycin B (Santa Cruz #sc-358688), tetracycline hydrochloride (Sigma-Aldrich #T7660).

### Cell culture and transfection

All cells utilized in this study were kept in Dulbecco′s Modified Eagle′s Medium - high glucose, DMEM (Sigma #D6429) supplemented with 1% Penicillin-Streptomycin (Sigma-Aldrich #P4333) and 10% FBS (Sigma-Aldrich, #F7524). The HeLa Flp-In T-REx cell line with EGFP-NBR1 integrated had their cell media supplied with 100 µg/ml hygromycin (Thermo Fisher Scientific #10687010) and 7.5 µg/ml Blasticidin (Thermofisher #A1113903). The NBR1 KO-p62 KO-, and NBR1/p62 double KO-(DKO) HeLa cell lines were maintained in media supplemented with 0.5 µg/ml puromycin (Sigma-Aldrich, P8833). All transfections were conducted in 24-well plates with cells at ∼50% confluency. TransIT®-LT1 Transfection Reagent (Mirus #MIR 2304) was used for transfection of DNA constructs. Leptomycin B (Santa Cruz #sc-358688) was used to inhibit nuclear export. To induce expression of EGFP-NBR1, HeLa Flp-In T-REx EGFP-NBR1 cells were treated with 1 µg/ml tetracycline hydrochloride (Sigma #T7660) for 24h.

### Generation of a stable cell line expressing EGFP-tagged NBR1

HeLa Flp-In T-REx cells were used to make a cell line stably expressing EGFP-tagged NBR1 from a tetracycline inducible promoter. Following the Flp-In system manual (Invitrogen), cDNA5/FRT/TO-EGFP-NBR1 were co-transfected with the Flp recombinase encoding plasmid pOG44 (Invitrogen, V6520-20) into the Flp-In T-REx HeLa host cell line. This cell line has a single chromosomal FRT site for insertion of selected cDNA constructs and expresses the TetR repressor enabling tetracycline regulated expression of inserted constructs. 48 hours after transfection, colonies of cells with EGFP-NBR1 integrated were selected with 200 ng/ml of hygromycin (Calbiochem, 400051).

### Generation of knockout cell lines sing CRISPR/Cas9

CRISPR/cas9-mediated knockout were generated as described (Abudu et al., 2021). Small guide RNA (sgRNA) designed to target different exons of p62 and NBR1 were annealed and ligated into *Bbs*1 linearized vectors carrying a wild-type CRISPR-associated protein 9 (Cas9) and either green fluorescent protein (GFP) (Addgene, #48138) or puromycin resistance gene (Addgene, #62988). HeLa cells seeded into 6cm plates were transfected with the sgRNA-containing Cas9 vectors using Metafectene Pro (Biontex #T040). 18-24h after transfection, cells were selected by either treatment with 1-3µg/ml puromycin for 24-36hrs for vectors carrying the puromycin gene or directly sorted by fluorescence-activated cell sorting using the EGFP signal. Single clones sorted into 96-well plates were then expanded. Gene knockouts were screened by immunoblotting and/or genomic analysis. For genomic screening, DNA was first extracted from cells using the GenElute mammalian genomic DNA miniprep kit (Sigma #G1N350) and the gene area of interest amplified by polymerase chain reaction (PCR). The PCR-amplified region was ligated into the pGEM-T vector (Promega #A3600) and sequenced to identify insertions or deletions (indels). CRISPR gRNA primers used include NBR1, GCCAGAGGATCCTGCAGTGC; p62, GGCGCCTCCTGAGCACACGG.

### Immunofluorescence

Cells were seeded on glass coverslips (VWR #631-0150) in 24-well plates. After the indicated treatment, cells were fixed in 4% PFA in PBS (137 mM NaCl, 2.7 mM KCl, 4.3 mM Na_2_HPO_4_, 1.47 mM KH_2_PO_4_, pH 7.4.) for 15 min at RT. The cells were then washed 3x in PBS and permeabilized with 0.2% Triton X-100 for 5 min. After permeabilization, the cells were washed 3x in PBS, followed by blocking in 2.5% goat serum in PBS for 1 h at RT. The cells were then incubated with primary antibodies overnight at 4°C, using the following dilutions: anti-p62/SQSTM1 (1:1000), anti-NBR1 (1:500), anti-lamin B1 (1:1000), anti-FK2 (1:1000), anti-PML (1:1000), anti-proteasome (1:25). Cells were incubated with Alexa fluor secondary antibodies (1:1000) for 1 h at RT after washing 3x in PBS. The cells were then washed an additional 3x in PBS prior to mounting of the coverslips to glass slides with ProLong™ Diamond Antifade Mountant (Thermo Fisher Scientific #P36965). The cells were examined with a LSM800 confocal laser microscope equipped with a Plan-Apochromat 63×/1.4 Oil DIC M27 objective (Zeiss).

### Fluorescence Recovery After Photobleaching (FRAP)

FRAP analysis was performed on live cells 48 h following transfection with EGFP-p62-ΔNES(Δ303-320), mCherry-NBR1-ΔNES1(Δ169-186)-ΔNES2(Δ557-574), EGFP-NBR1-D50R-ΔNES1ΔNES2, or a combination of these constructs. A Zeiss LSM800 confocal microscope equipped with a 63x/1.4 NA oil immersion objective was used, together with an incubation chamber (5% CO_2_, 37°C). Individual nuclear foci were bleached using 2 µm circular regions of interest (ROI), using a single pulse of the 488 nm laser at 80% power. An unbleached control ROI was also monitored, with a maximum of 2 bleached ROI’s per nucleus. Images were captured at 1 sec intervals using 1% power, with 1 AU pinhole, and using the 4x optical zoom setting. Recovery and half-time curves were generated by using ZEN Blue (Zeiss) and Graphpad Prism software, after correcting for photobleaching and scaling values between 0 (post-bleach value) and 1 (pre-bleach value). Dots that did not recover were excluded from the dataset.

### 1,6-hexanediol studies

For 1,6-hexanediol (1,6-HD) experiments, a 3.5% solution was made by diluting 1,6-HD (Sigma-Aldrich, #240117) in a 1:3 ratio of complete medium to sterile distilled water. Cells were seeded in 8-well chamber dishes and transfected as for FRAP experiments. and imaged using an incubation chamber (37°C, 5% CO_2_). Control images were captured before replacing the cell medium with the pre-warmed 1,6-HD solution. Images were immediately captured and subsequently at 1 min intervals to analyze short-term effects. Condensate recovery was assessed by removing the 1,6-HD solution from wells and replacing it with fresh pre-warmed medium. Images were captured after 30 and 60 min.

### Western blot experiments

Cells were washed in PBS followed by lysis directly in SDS-PAGE loading buffer (2% SDS, 10% glycerol 50 mM Tris-HCl, pH 7.4) supplemented with 200 mM dithiothreitol (DTT, Sigma, #D0632). Lysates were heated at 100° C for 10 min. Protein concentration was measured and bromophenol blue (0.1%) added. Samples (10-60 µg) were run on 10% SDS-polyacrylamide gels and blotted on Hybond nitrocellulose membranes (GE Healthcare, 10600003) followed by Ponceau S staining. Blocking was done in 5% nonfat dry milk in 1% TBS-T (0.2M Tris-HCl, pH 8.0, 1.5 M NaCl, 0.05% Tween-20 (Sigma, #P9416)). The primary antibody was diluted in PBS-Tween 20 containing 5% nonfat dry milk before incubation for 24h at 4°C. Membranes were washed 3 times (10 min each) with TBS-T. Incubation with secondary antibody was done at room temperature for 1 h in PBS-Tween 20 containing 5% nonfat dry milk. Membranes were washed 3 times (10 min each) and analyzed by enhanced chemiluminescence using ImageQuant LAS-4000 (GE Healthcare).

### Bioinformatics and statistics

Post acquisition analysis of images was done with ImageJ and custom-made scripts for quantification. Statistical parameters are denoted in the figure legends. To determine statistical significance, one-way ANOVA and two-way ANOVA were used, with a P value of <0.05 considered significant. Tukey’s multiple comparisons test was applied post hoc to assess the differences between groups.

## Acknowledgements

We are grateful to the proteomics and imaging core facilities at UiT, Faculty of Health Sciences, for valuable assistance.

## Author contributions

Conceptualization: M.M.M., T.J., T.L.; Methodology: M.M.M., J.W., M.S.N., G.E., Y.P.A., H.L.O.; Validation: M.M.M., J.W., T.J., T.L.; Formal analysis: M.M.M., J.W., M.S.N.; Investigation: M.M.M., J.W., M.S.N., H.L.O., T.L., G.E.; Resources: T.L., G.E., Y.P.A,; Writing - original draft: M.M.M., J.W., T.L, T.J.; Writing - review & editing: T.J., T.L., H.L.O; Visualization: M.M.M., J.W., T.J., T.L.; Supervision: T.J., T.L.; Project administration: T.L, T.J.; Funding acquisition: T.J.

---

**Figure S1.**
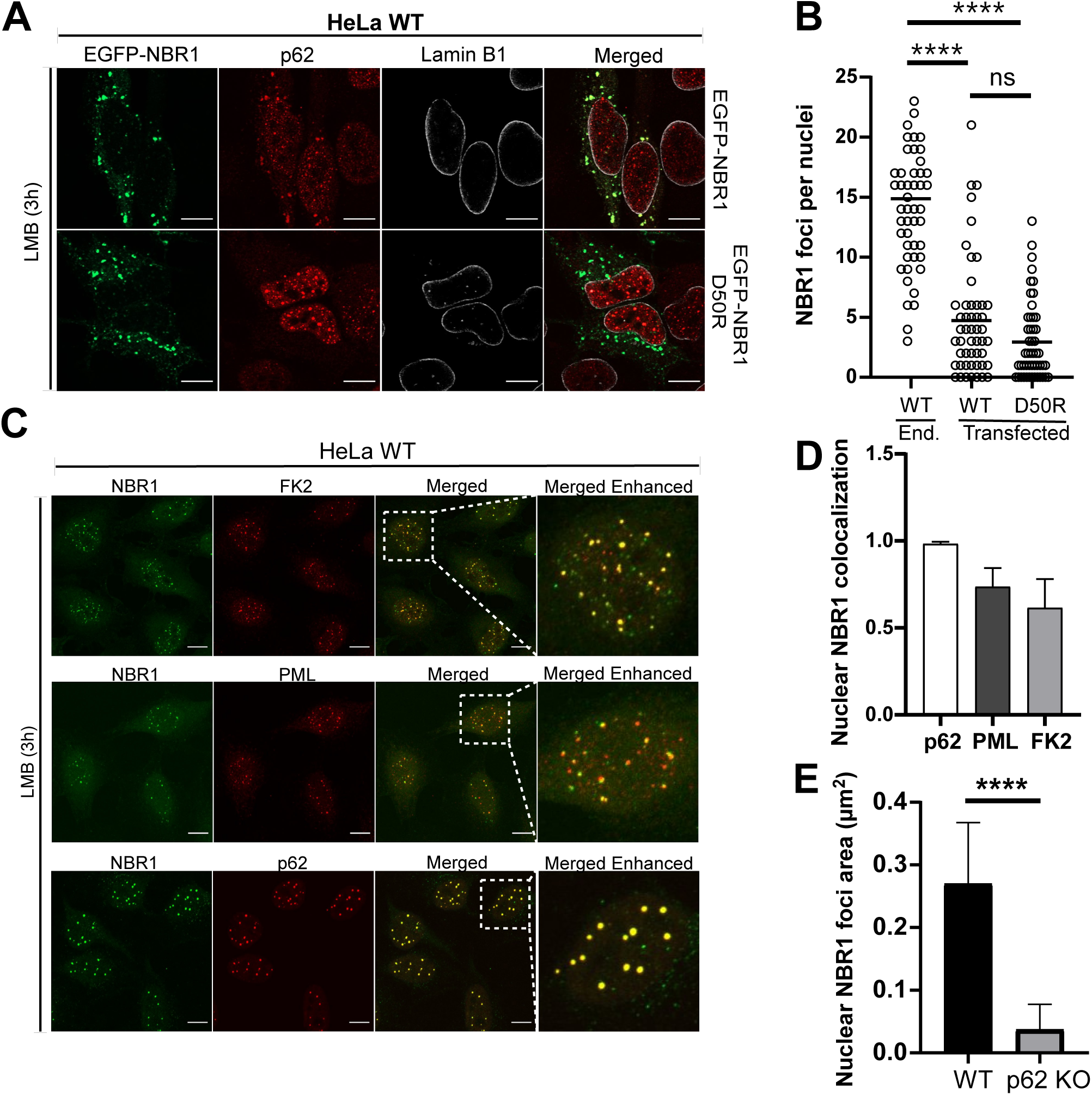
Endogenous NBR1 accumulates in nuclear p62 bodies. (A) HeLa cells transfected with EGFP-NBR1 or EGFP-NBR1 D50R (green). Cells were stained for NBR1 (green), p62 (red) and lamin B1 (white). (B) Quantification of data in A: Nuclear NBR1 foci per cell (n=50 cells). Statistical comparison by One-way ANOVA. ****p<0.0001. (C) HeLa cells were treated with LMB (3 h) and stained for NBR1 (green) and one of the following (red): FK2 (ubiquitylated proteins), PML (promyelocytic leukemia protein), or p62. Scale bars equal to 10 µm. (D) Quantification of colocalization of nuclear NBR1 puncta with FK2, PML, or p62. Data represent mean ± SD (n=50 cells). (E) Quantification of mean ± SD area of NBR1 puncta (µm²) (n=50 cells). Statistical comparison by One-way ANOVA. ****p < 0.0001.

**Figure S2.**
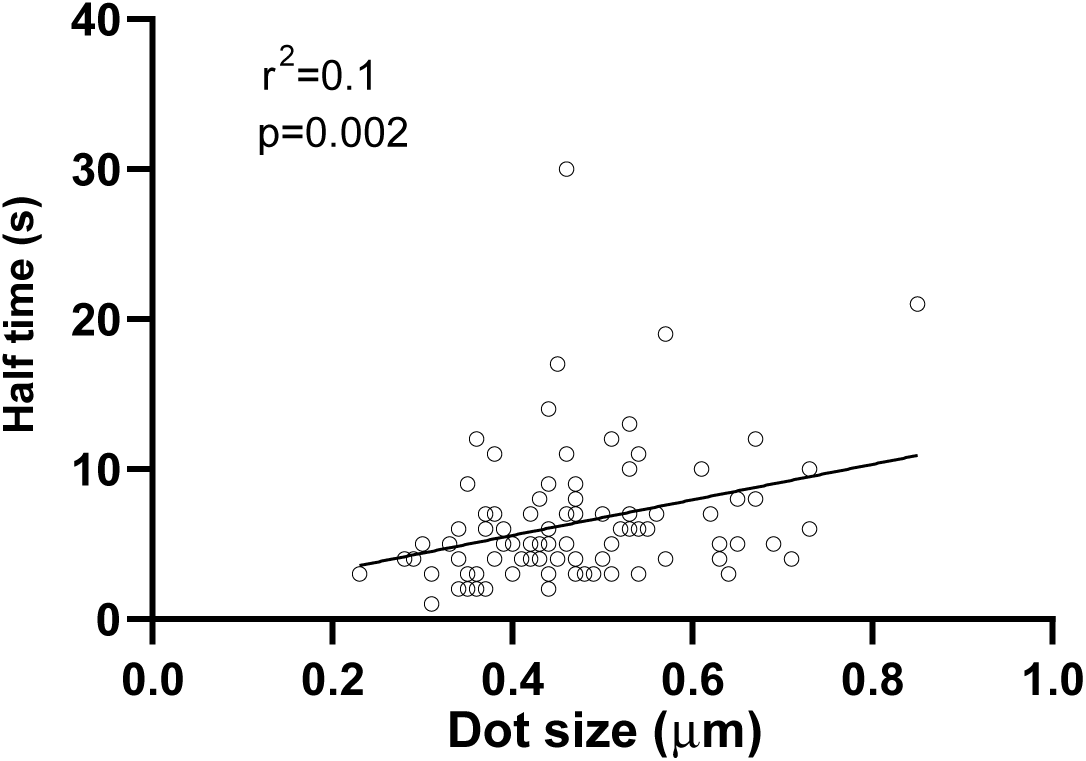
In FRAP experiments the recovery rate may depend on the size of nuclear puncta. FRAP analyses of EGFP positive puncta in HeLa p62/NBR1 DKO cells transfected with EGFP-p62ΔNES, indicating an effect of puncta size on recovery rate.

## REFERENCES

Abudu, Y.P., S. Mouilleron, S.A. Tooze, T. Lamark, and T. Johansen. 2021. SAMM50 is a receptor for basal piecemeal mitophagy and acts with SQSTM1/p62 in OXPHOS-induced mitophagy. Autophagy. 17:2656–2658.

Bjørkøy, G., T. Lamark, A. Brech, H. Outzen, M. Perander, A. Øvervatn, H. Stenmark, and T. Johansen. 2005. p62/SQSTM1 forms protein aggregates degraded by autophagy and has a protective effect on huntingtin-induced cell death. J Cell Biol. 171:603–614.

Ciuffa, R., T. Lamark, A.K. Tarafder, A. Guesdon, S. Rybina, W.J. Hagen, T. Johansen, and C. Sachse. 2015. The Selective Autophagy Receptor p62 Forms a Flexible Filamentous Helical Scaffold. Cell Rep. 11:748–758.

Cummings, C.J., M.A. Mancini, B. Antalffy, D.B. DeFranco, H.T. Orr, and H.Y. Zoghbi. 1998. Chaperone suppression of aggregation and altered subcellular proteasome localization imply protein misfolding in SCA1. Nat Genet. 19:148–154.

Deosaran, E., K.B. Larsen, R. Hua, G. Sargent, Y. Wang, S. Kim, T. Lamark, M. Jauregui, K. Law, J. Lippincott-Schwartz, A. Brech, T. Johansen, and P.K. Kim. 2013. NBR1 acts as an autophagy receptor for peroxisomes. J Cell Sci. 126:939–952.

Ernoult-Lange, M., A. Wilczynska, M. Harper, C. Aigueperse, F. Dautry, M. Kress, and D. Weil. 2009. Nucleocytoplasmic traffic of CPEB1 and accumulation in Crm1 nucleolar bodies. Mol Biol Cell. 20:176–187.

Fu, A., V. Cohen-Kaplan, N. Avni, I. Livneh, and A. Ciechanover. 2021. p62-containing, proteolytically active nuclear condensates, increase the efficiency of the ubiquitin-proteasome system. Proc Natl Acad Sci U S A. 118.

Fu, A., Z. Luo, T. Ziv, X. Bi, C. Lulu-Shimron, V. Cohen-Kaplan, and A. Ciechanover. 2024. Nuclear p62 condensates stabilize the promyelocytic leukemia nuclear bodies by sequestering their ubiquitin ligase RNF4. Proc Natl Acad Sci U S A. 121:e2414377121.

Henderson, B.R., and A. Eleftheriou. 2000. A comparison of the activity, sequence specificity, and CRM1-dependence of different nuclear export signals. Exp Cell Res. 256:213–224.

Jakobi, A.J., S.T. Huber, S.A. Mortensen, S.W. Schultz, A. Palara, T. Kuhm, B.K. Shrestha, T. Lamark, W.J.H. Hagen, M. Wilmanns, T. Johansen, A. Brech, and C. Sachse. 2020. Structural basis of p62/SQSTM1 helical filaments and their role in cellular cargo uptake. Nat Commun. 11:440.

Kirkin, V., T. Lamark, Y.S. Sou, G. Bjorkoy, J.L. Nunn, J.A. Bruun, E. Shvets, D.G. McEwan, T.H. Clausen, P. Wild, I. Bilusic, J.P. Theurillat, A. Overvatn, T. Ishii, Z. Elazar, M. Komatsu, I. Dikic, and T. Johansen. 2009. A role for NBR1 in autophagosomal degradation of ubiquitinated substrates. Mol Cell. 33:505–516.

Kroschwald, S., S. Maharana, and A. Simon. 2017. Hexanediol: a chemical probe to investigate the material properties of membrane-less compartments. Matters.

Kudo, N., N. Matsumori, H. Taoka, D. Fujiwara, E.P. Schreiner, B. Wolff, M. Yoshida, and S. Horinouchi. 1999. Leptomycin B inactivates CRM1/exportin 1 by covalent modification at a cysteine residue in the central conserved region. Proc Natl Acad Sci U S A. 96:9112–9117.

Lamark, T., and T. Johansen. 2021. Mechanisms of Selective Autophagy. Annu Rev Cell Dev Biol. 37:143–169.

Lamark, T., M. Perander, H. Outzen, K. Kristiansen, A. Øvervatn, E. Michaelsen, G. Bjørkøy, and T. Johansen. 2003. Interaction codes within the family of mammalian Phox and Bem1p domain-containing proteins. J Biol Chem. 278:34568–34581.

Larsen, K.B., T. Lamark, A. Øvervatn, I. Harneshaug, T. Johansen, and G. Bjørkøy. 2010. A reporter cell system to monitor autophagy based on p62/SQSTM1. Autophagy. 6:784–793.

Lee, Y., J. Pei, J.M. Baumhardt, Y.M. Chook, and N.V. Grishin. 2019. Structural prerequisites for CRM1-dependent nuclear export signaling peptides: accessibility, adapting conformation, and the stability at the binding site. Sci Rep. 9:6627.

Lu, J., T. Wu, B. Zhang, S. Liu, W. Song, J. Qiao, and H. Ruan. 2021. Types of nuclear localization signals and mechanisms of protein import into the nucleus. Cell Commun Signal. 19:60.

Ma, J., T. Zhang, V. Novotny-Diermayr, A.L. Tan, and X. Cao. 2003. A novel sequence in the coiled-coil domain of Stat3 essential for its nuclear translocation. J Biol Chem. 278:29252–29260.

Mardakheh, F.K., G. Auciello, T.R. Dafforn, J.Z. Rappoport, and J.K. Heath. 2010. Nbr1 is a novel inhibitor of ligand-mediated receptor tyrosine kinase degradation. Mol Cell Biol. 30:5672–5685.

Matsumoto, G., K. Wada, M. Okuno, M. Kurosawa, and N. Nukina. 2011. Serine 403 Phosphorylation of p62/SQSTM1 Regulates Selective Autophagic Clearance of Ubiquitinated Proteins. Mol Cell. 44:279–289.

Nagaoka, U., K. Kim, N.R. Jana, H. Doi, M. Maruyama, K. Mitsui, F. Oyama, and N. Nukina. 2004. Increased expression of p62 in expanded polyglutamine-expressing cells and its association with polyglutamine inclusions. J Neurochem. 91:57–68.

Neufeld, K.L., D.A. Nix, H. Bogerd, Y. Kang, M.C. Beckerle, B.R. Cullen, and R.L. White. 2000. Adenomatous polyposis coli protein contains two nuclear export signals and shuttles between the nucleus and cytoplasm. Proc Natl Acad Sci U S A. 97:12085–12090.

Nguyen Ba, A.N., A. Pogoutse, N. Provart, and A.M. Moses. 2009. NLStradamus: a simple Hidden Markov Model for nuclear localization signal prediction. BMC Bioinformatics. 10:202.

Pankiv, S., T. Lamark, J.A. Bruun, A. Øvervatn, G. Bjørkøy, and T. Johansen. 2010. Nucleocytoplasmic shuttling of p62/SQSTM1 and its role in recruitment of nuclear polyubiquitinated proteins to promyelocytic leukemia bodies. J Biol Chem. 285:5941–5953.

Pikkarainen, M., P. Hartikainen, and I. Alafuzoff. 2008. Neuropathologic features of frontotemporal lobar degeneration with ubiquitin-positive inclusions visualized with ubiquitin-binding protein p62 immunohistochemistry. J Neuropathol Exp Neurol. 67:280–298.

Rasmussen, N.L., A. Kournoutis, T. Lamark, and T. Johansen. 2022. NBR1: The archetypal selective autophagy receptor. J Cell Biol. 221.

Sanchez-Martin, P., Y.S. Sou, S. Kageyama, M. Koike, S. Waguri, and M. Komatsu. 2020. NBR1-mediated p62-liquid droplets enhance the Keap1-Nrf2 system. EMBO Rep. 21:e48902.

Sheldon, L.A., and R.E. Kingston. 1993. Hydrophobic coiled-coil domains regulate the subcellular localization of human heat shock factor 2. Genes Dev. 7:1549–1558.

Shin, H.Y., and N.C. Reich. 2013. Dynamic trafficking of STAT5 depends on an unconventional nuclear localization signal. J Cell Sci. 126:3333–3343.

Souquere, S., D. Weil, and G. Pierron. 2015. Comparative ultrastructure of CRM1-Nucleolar bodies (CNoBs), Intranucleolar bodies (INBs) and hybrid PML/p62 bodies uncovers new facets of nuclear body dynamic and diversity. Nucleus. 6:326–338.

Terasawa, H., Y. Noda, T. Ito, H. Hatanaka, S. Ichikawa, K. Ogura, H. Sumimoto, and F. Inagaki. 2001. Structure and ligand recognition of the PB1 domain: a novel protein module binding to the PC motif. EMBO J. 20:3947–3956.

Turco, E., A. Savova, F. Gere, L. Ferrari, J. Romanov, M. Schuschnig, and S. Martens. 2021. Reconstitution defines the roles of p62, NBR1 and TAX1BP1 in ubiquitin condensate formation and autophagy initiation. Nat Commun. 12:5212.

Ura, S., N. Masuyama, J.D. Graves, and Y. Gotoh. 2001. Caspase cleavage of MST1 promotes nuclear translocation and chromatin condensation. Proc Natl Acad Sci U S A. 98:10148–10153.

Wilson, M.I., D.J. Gill, O. Perisic, M.T. Quinn, and R.L. Williams. 2003. PB1 domain-mediated heterodimerization in NADPH oxidase and signaling complexes of atypical protein kinase C with Par6 and p62. Molecular cell. 12:39–50.

Xu, D., K. Marquis, J. Pei, S.C. Fu, T. Cagatay, N.V. Grishin, and Y.M. Chook. 2015. LocNES: a computational tool for locating classical NESs in CRM1 cargo proteins. Bioinformatics. 31:1357–1365.

You, Z., W.X. Jiang, L.Y. Qin, Z. Gong, W. Wan, J. Li, Y. Wang, H. Zhang, C. Peng, T. Zhou, C. Tang, and W. Liu. 2019. Requirement for p62 acetylation in the aggregation of ubiquitylated proteins under nutrient stress. Nat Commun. 10:5792.

Zaffagnini, G., A. Savova, A. Danieli, J. Romanov, S. Tremel, M. Ebner, T. Peterbauer, M. Sztacho, R. Trapannone, A.K. Tarafder, C. Sachse, and S. Martens. 2018. p62 filaments capture and present ubiquitinated cargos for autophagy. EMBO J. 37.

